# A recently collected *Xanthomonas translucens* isolate encodes TAL effectors distinct from older, less virulent isolates

**DOI:** 10.1101/2023.09.15.558010

**Authors:** Diego E. Gutierrez-Castillo, Emma Barrett, Robyn Roberts

## Abstract

*Xanthomonas translucens,* the causal agent of bacterial leaf streak disease (BLS) in cereals, is a re-emerging pathogen that is becoming increasingly destructive across the world. While BLS has caused yield losses in the past, there is anecdotal evidence that newer isolates may be more virulent. We observed that two *Xanthomonas translucens* isolates collected from two sites in Colorado are more aggressive on current wheat and barley varieties compared to older isolates, and we hypothesize that genetic changes between recent and older isolates contribute to the differences in isolate aggressiveness. To test this, we phenotyped and genetically characterized two *X. translucens* isolates collected from Colorado in 2018, which we designated CO236 (from barley) and CO237 (from wheat). Using pathovar-specific phenotyping and PCR primers, we determined that CO236 belongs to pathovar translucens and CO237 belongs to pathovar undulosa. We sequenced the full genomes of the isolates using Oxford Nanopore long-read sequencing, and compared their whole genomes against published *X. translucens* genomes. This analysis confirmed our pathovar designations for Xtt CO236 and Xtu CO237, and showed that, at the whole-genome level, there were no obvious genomic structural changes between Xtt CO236 and Xtu CO237 and other respective published pathovar genomes. Focusing on pathovar undulosa (Xtu CO237), we then compared putative Type III effectors among all available Xtu isolate genomes and found that they were highly conserved. However, there were striking differences in the presence and sequence of various transcription activator-like effectors (TALE) between Xtu CO237 and published undulosa genomes, which correlate with isolate virulence. Here, we explore the potential implications of the differences in these virulence factors, and provide possible explanations for the increased virulence of recently-emerged isolates.

**Data Summary:** A list of all *Xanthomonas* accessions used in this study can be found in Supplemental Table S1. Xtt CO236 and Xtu CO237 genomic sequences are deposited in GenBank (Accession: PRJNA1017868 and PRJNA1017870, respectively). Software packages for the custom Conda environment used in this analysis can be found in Supplemental Table S4. The dataset from the MinION reads from CO236 and CO237 can be found in Dryad, https://doi.org/10.5061/dryad.d51c5b06q. Custom bash and Python scripts for the effector analysis are available in (https://github.com/robertslabcsu/xanthanalysis.git).

**Impact statement:** *Xanthomonas translucens* is a destructive, re-emerging pathogen of cereal crops with no known resistance or methods for chemical control. Recent isolates have increased virulence compared to older isolates, which emphasizes the need to understand how virulence evolves, and how the pathogen interacts with its host, to find new ways to manage the disease. Here, we identify potential virulence factors that contribute to the increased aggressiveness observed in two recently collected Colorado isolates, with potential impacts on understanding pathogen host range and evolution.

## INTRODUCTION

Wheat is one of the most important cereals globally, with 431 million tons produced worldwide in 2021, and 30 million tons from the USA alone [1]. Wheat is the 3^rd^ largest acreage under cultivation in the US, following corn and soybeans. Of the diseases of wheat, bacterial leaf streak (BLS), caused by *Xanthomonas translucens* (Xt), is a major concern and also infects many other cereals including barley, rye, and wild grasses [2, 3]. Yield losses due to Xt are difficult to measure, but a recent study suggests that yield losses may be up to 60%, especially since all varieties are susceptible [4]. There is no commercial resistance available, and no chemicals can be used to treat or prevent the disease. Moreover, little is known about the molecular mechanisms behind wheat immune responses, and even less is known for BLS disease.

There are eight different pathovars of *X. translucens* [3], but the most economically impactful to cereal crops are pathovars undulosa and translucens. These pathovar designations are based on nomenclature for species with distinguishable phenotype and host range [5]. The *X. translucens* pathovar undulosa (Xtu) primarily infects wheat and causes some disease on barley, while the *X. translucens* pathovar translucens (Xtt) primarily infects barley and generally causes little disease on wheat. Strains associated with specific pathovars also have distinct sequence types [6]. Similarly, *X. translucens* pathotype strains group into three genetically distinct clades within the species [7]. Biologically, Xtu is thought to primarily move through the leaf apoplast, whereas Xtt likely moves primarily through the leaf vasculature [8]. Recently, using several publicly available genomes, primers were designed to distinguish undulosa and translucens from other Xt pathovars based on genes predicted to be specific for pathogen infection and establishment mechanisms [9].

Nearly 80 Xt genomes are publicly available, and many of these have been assembled using long-read sequencing, including PacBio and Oxford Nanopore Technology (ONT), which may or may not be corrected with short-read sequencing such as Illumina [10–12]. While long-read sequencing is useful in detecting large polymorphisms or structural variation, the base sequence accuracy is lower than with short-read sequencing [13]. Genomic ‘polishing’ tools have been developed to increase accuracy using various algorithms [14], which have been designed especially for Nanopore sequencing, including tools such as Medaka [15] and Homopolish [16]. In particular, genomic regions high in repetitive sequence can be difficult to assemble strictly via long-read sequencing. *Xanthomonas* genomes often contain regions of high repetition, and tools using computational corrections have been designed even for specific gene classes such as transcription activator-like effectors (TALEs), which are important virulence factors for many *Xanthomonas* species [17].

Virulence factors aid bacteria and other pathogens in infection of hosts. Often, virulence factors are proteins, called effectors, that either increase plant susceptibility by making nutrients more available to the pathogen, or by suppressing the host immune system and related pathways. The type III effectors (T3Es) are proteins that are delivered into hosts through a needle-like apparatus, the type III secretion system (T3SS). These T3Es contribute to bacterial pathogenesis by suppressing the basal plant defense responses, modulating MAPK cascades involved in defense, interfering with plant immunity, regulating hormones, modulating host gene expression, and disrupting the plant cytoskeleton [18]. One class of T3Es specific to *Xanthomonas* are the TALEs, which are expressed by some, but not all, *Xanthomonas* species. Other bacterial species have TALE orthologs with similar putative ontology, including strains from the *Ralstonia solanacearum* species complex (RipTALs) [19] and *Burkholderia* (BTLs) [20]. To study the high diversity of TALEs across strains from the same species and across the genus, a hierarchical and agglomerative clustering approach that groups TALEs based on their repeated variable diresidue (RVD) sequence is used [21]. The classes derived from this approach have nomenclature that begins with ‘Tal’ and is followed by a unique two-character identifier, with each individual TALE labeled with a number to identify it within the class (e.g TalAD1). Although BLS disease has become more prevalent across the world, still very few reports exist on the known virulence factors and effectors of *X. translucens* in cereals. Two examples are of a TAL effector in Xtu 4699 (Tal8) that has been characterized for its role in commandeering a host-limiting step in abscisic acid biosynthesis [22], and Tal1 in a *X. translucens* pv. cerealis (Xtc01) that contributes to virulence in wheat [23].

Here, we investigated two recently collected and highly virulent Xt isolates found in Colorado in 2018. We characterized these isolates for their virulence and determined the pathovar classifications by phenotype and genotype. Using whole-genome sequencing, we compared these isolates with each other and to other publicly available Xt genomes to find gene candidates that may be strong contributors to virulence of the Colorado Xtu isolate.

## METHODS

### Strain and pathovar confirmation

Two *Xanthomonas translucens* isolates were collected in 2018 in Colorado from barley and wheat with bacterial leaf streak (BLS) disease. Isolate CO236 was collected from barley in the San Luis Valley, and isolate CO237 from wheat in Akron. The isolates were streaked to single colonies, then grown at 28°C in yeast dextrose carbonate agar medium (1% yeast extract, 2% dextrose, 2% calcium carbonate, 1.5% agar). Observed was the typical yellow and mucoid phenotype of *Xanthomonas* strains [24]. To confirm CO236 and CO237 pathovar classification, the isolates were grown on nutrient agar, and then the bacteria were suspended in 10mL of 10mM MgCl_2_. Bacteria were pelleted at 4500 RPM for 7 min. The supernatant was removed, and the bacterial pellet was resuspended in 10mM MgCl_2_ to wash the pellet. The final bacterial pellet was resuspended in 5mL of 10mM MgCl_2_. 200µL of the bacterial suspension was used to extract genomic DNA (gDNA) using the Zymo Research Quick-DNA™ Fungal/Bacterial Miniprep Kit, according to the manufacturer’s instructions.

The pathovar group of isolates CO236 and CO237 was determined using multiplex PCR, with comparisons to a known Xtt UPB886 [25], Xtu ICMP1105 [26] and *X. vasicola* pv. *vasculorum* NE-433 [27]. The multiplex PCR for this test consisted of a 25µL reaction with primers *cbsA-1*, *cbsA-2*, *cbsA-3*, *cbsA-4* at a concentration of 0.08µM each [25], the pair of S8-protease primers (0.16µM each), 5µL of 5X Q5 reaction buffer with Q5 High G/C enhancer, 0.5 U of Q5 High-Fidelity DNA Polymerase, 0.5µL of 10mM dNTP mix, 2µL of genomic DNA, and nuclease-free water up to 25µL. The conditions for the PCR were an initial denaturation at 98°C for 2 min, 30 cycles at 98°C for 30 sec, 50°C for 45 sec, and 72°C for 30 sec, and a final extension at 72°C for 2 min. The PCR products were visualized in a 0.8% agarose gel stained with GelRed^TM^ (GoldBio) using 80V for 40 minutes and visualized under blue light with a Gel Imager C300 (Azure biosystems).

### Bacterial inoculation and growth curves

After confirming their phenotypes *in vitro*, we grew the isolates on nutrient agar (0.5% peptone, 0.3% yeast extract, 1.5% agar, 0.5% NaCl) for 48 hours for wheat (var. Hatcher) and barley (var. Morex) inoculations to confirm their phenotypes *in planta*. The Xtu isolates were suspended in MgCl_2_ as described previously [7] and adjusted to a final OD of 0.001 (∼1.0 x 10^6^ colony forming units/mL) after two consecutive washes in the same buffer. These dilutions were used to inoculate whole leaves of wheat and barley making 4 and 5 infiltrations respectively to ensure inoculation of the whole area of the leaf. Bacterial populations were measured 0, 48 and 96 hpi, from six leaf discs obtained from the infiltrated area using a 0.1cm diameter cork-borer. The discs were immediately placed in 2mL tubes with two metal beads, and flash-frozen on liquid nitrogen. Samples were stored at −80°C until they were ready to be processed. The samples were ground in a Tissue Lyser II (Qiagen), then suspended in 0.4mL of sterile distilled water. The bacterial suspensions were diluted and 20µL per sample was plated on nutrient agar to count colonies. Dilutions yielding 25-300 colony forming units (CFU) were chosen to calculate the total number of CFU/cm^2^ for each sample. The Xtt isolates were pre-treated similarly as described above and prepared to a final OD of 0.4 (∼4.0 x 10^8^ CFU/mL) in 50mL conical tubes. Scissors disinfected with 75% ethanol were dipped and opened/closed inside the bacterial suspension. Immediately after, the soaked scissors were used to clip the 2^nd^ or 3^rd^ leaf. Between strains, scissors were disinfected with 75% ethanol. Plants were evaluated for 14 days post clip inoculation to measure lesion length.

### Genomic DNA extraction and Whole-genome sequencing

The gDNA obtained was checked for quality by running on an agarose gel (1%), and we confirmed that no product smaller than 5kb was detected. Approximately 300 ng of gDNA from strains Xtt CO236 and Xtu CO237 was barcoded with Oxford Nanopore Technologies Rapid Barcoding Kit (SQK-RBK004). Barcoded libraries were prepared for Oxford Nanopore sequencing using the Rapid Sequencing Kit (SQK-RAD004), following the manufacturer’s instructions for whole-genome sequencing. The sequencing was carried out in a R 9.4.1 flow cell for 96 h and yielded raw read data consisting of 8.9 Gb of base-called reads for Xtt CO236, and 12 Gb for Xtu CO237. Base calling was done in real time with Guppy’s high-accurate algorithm from Oxford Nanopore Technologies.

### De Novo Genome Assembly and manual curation of TAL effector

The full bioinformatic pipeline was carried out on the Alpine supercomputer [28]. The assembly was carried out in a custom Conda 4.7.10 environment. The output from Guppy’s basecalled reads was filtered using Filtlong 0.2.1[29, 30] to discard low-quality reads, and the upper 90^th^ percentile with the highest quality score was used to assemble a draft genome with Flye 2.9 [31]. The draft genome was polished twice with the raw fastq reads using Medaka [15]. The full list of packages can be found in Supplemental Table 5. Since the polishing steps did not correct homopolymers commonly found in ONT-only assemblies on TAL-effector regions, a correction step was performed using Homopolish [32]. On the final assembly, TAL effectors were annotated with AnnoTALE [21]. The curated assemblies were formatted into GenBank files and uploaded into NCBI under the accession numbers PRJNA1017868 (Xtt CO236) and PRJNA1017870 (Xtu CO237).

### Effector Analysis across the undulosa pathovar

A custom bash script (https://github.com/robertslabcsu/xanthanalysis.git) was used to compare previously characterized *Xanthomonas* effectors against putative proteins of all NCBI-available (as of August 2023) Xtu genomes predicted by Prokka 1.13 [33] with tblastn. The custom list of effectors was extracted from the *Xanthomonas* Resource database (http://www.biopred.net/xanthomonas/t3e.html). The list used for the analysis can be found in Supplemental Table 4.

## RESULTS

### Two *Xanthomonas translucens* isolates from Colorado cause severe disease in wheat and barley

*Xanthomonas translucens* causes occasional disease epidemics in Colorado, and in 2018 there was an unusually severe outbreak. Two isolates were collected from leaves with apparent leaf streak disease: one from barley located in the San Luis Valley, CO (designated CO236), and one from wheat located in Akron, CO (designated CO237). Bacteria were isolated from the leaves and plated on yeast extract-dextrose-CaCO_3_ (YDC) medium, and single colonies were grown in culture and inoculated into wheat and barley by syringe infiltration. Symptoms were consistent with leaf streak disease produced by other *X. translucens* isolates from our collection (**Figure 1A)**, including water soaking, yellowing, and lesion streaks running parallel to the leaf veins. To determine which pathovars these isolates represented, bacteria were syringe-infiltrated at a concentration of 10^6^ CFU/ml into wheat (variety Hatcher) and barley (variety Morex) to determine host specificity (**Figure 1A**). The translucens pathovar is typically vascular and mainly infects barley, whereas the undulosa pathovar is primarily apoplastic and mostly infects wheat, though it can cause lighter water-soaking symptoms in barley [3]. We observed that CO236 symptoms were consistent with *Xt* pv. translucens (Xtt), and CO237 symptoms were consistent with *Xt* pv. undulosa (Xtu). To further confirm the identities of CO236 and CO237, *Xt* pathovar-specific primers amplifying the *cbsA* gene [9] were used in PCR on isolate genomic DNA (**Figure 1B**). Amplicons matched the expected sizes of *Xt* pv. translucens (CO236) and *Xt.* pv. undulosa (CO237). A negative control (*X. vesicatoria* pv. vesicatoria NE-433, Xvv) was used as an outgroup, which amplifies a different band size and confirms pathovar specificity for translucens and undulosa.

**Figure 1.**
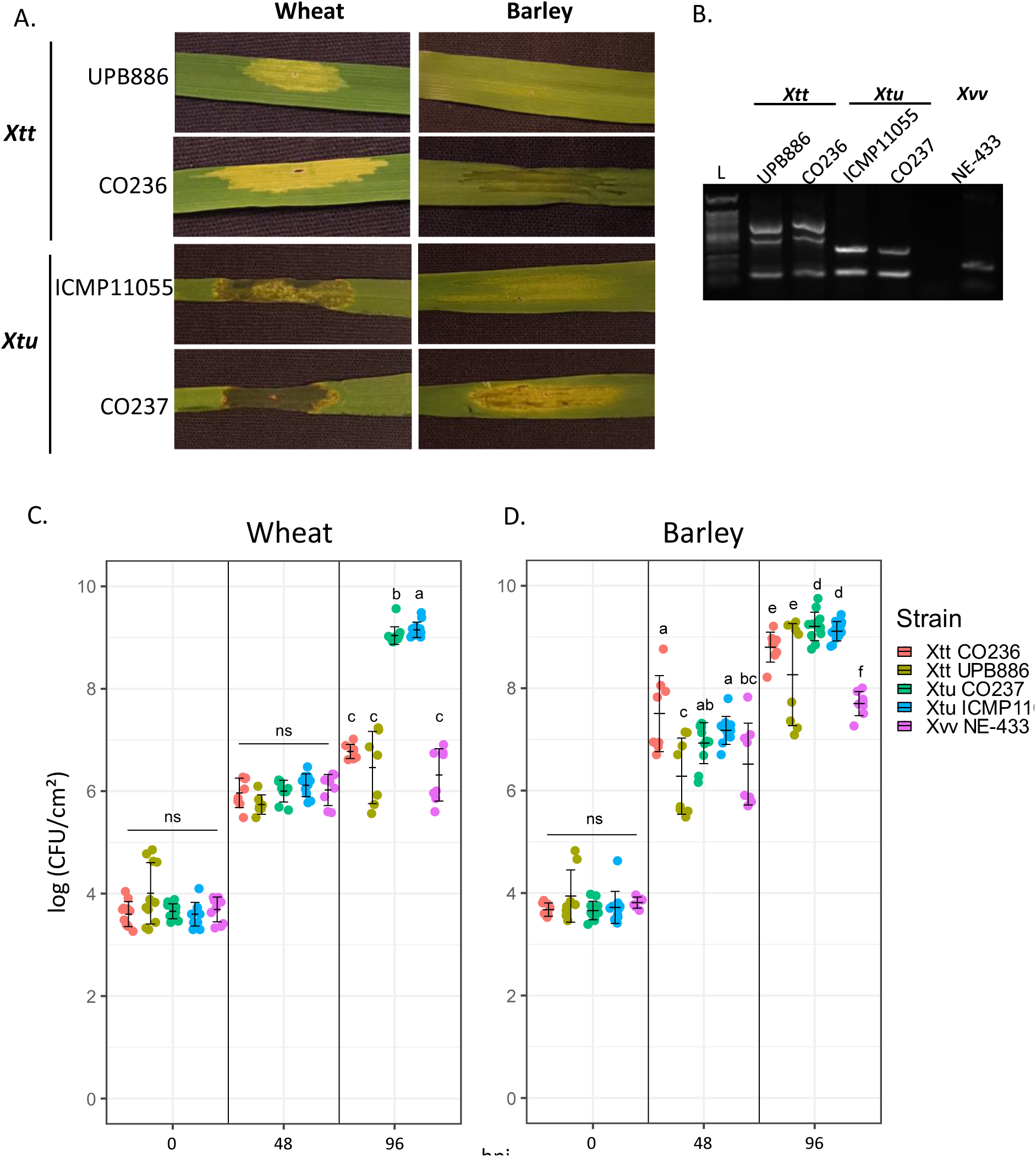
Phenotypic and molecular characterization of *Xanthomonas translucens* isolates from Colorado. A. Symptoms of wheat (variety Hatcher) and barley (variety Morex) leaves after syringe-infiltration of Xtt and Xtu isolates (1×10^6^ CFU/mL). Virulence of Colorado isolates (Xtt CO236, Xtu CO237) was compared with isolates UPB886 (Xtt, 1990, Iran) and ICMP11055 (Xtu, 1983, Iran). **B.** Isolate characterization using pathovar-specific *X.t.* pv. undulosa and translucens primers (Roman-Reyna et al., 2020). Xtt UPB886 and Xtu ICMP11055 were used as positive controls, while Xvv NE-433 was used as an unrelated *Xanthomonas* pathogen control. L, ladder. **C.** Bacterial populations of CO237, ICMP11055 and NE-433 at 0, 24, and 48 hours post syringe-inoculation (hpi) in wheat leaves. **D.** Bacterial populations of CO237, ICMP11055 and NE-433 at 0, 24, and 48 h post syringe-inoculation in barley leaves. Significance determined via Wilcoxon test in R software (p < 0.05). N= 12.

We also observed that, compared to ICMP11055, an aggressive reference isolate of Xtu, CO237 was more virulent and caused more necrosis and water-soaking symptoms on both wheat and barley when inoculated by syringe infiltration. Xtt CO236 also was more virulent than the Xtt reference UPB886, also highly virulent, causing more water-soaking in barley and more extensive and intense yellowing in wheat. Because Xtt is generally considered a vascular pathogen, we clip-inoculated both CO236 and UPB886 into barley to see if symptoms would be different depending on inoculation method (**Supplemental Figure S1**). We found no significant difference in lesion lengths between Xtt CO236 and Xtt UPB886, suggesting that inoculation method does not impact differences in Xtt bacterial movement. To determine if the increased virulence of CO237 and CO236 was associated with increased bacterial populations, we syringe-infiltrated leaves with a bacterial suspension (OD_600_ = 0.001) in wheat and barley, and counted bacterial populations at 0-, 24-, and 48-hours post infiltration (hpi). Bacterial numbers were calculated as CFU/cm^2^ to standardize the infiltrated area of the leaf and compare populations between hosts (wheat and barley). Xvv, which is a maize pathogen, was used as a non-host pathovar control (**Figure 1C**). In wheat, we found no significant difference in bacterial populations at 0- and 48-hpi. At 96hpi, the Xtt strains (CO236 and UPB886) were not significantly different from the Xvv non-host control. Both Xtu strains grew significantly more than Xvv, and Xtu ICMP11055 populations were slightly higher than Xtu CO237. In barley, the Xtt and Xtu strains grew more than 10-fold over the Xvv non-host control. Surprisingly, both Xtu strains multiplied significantly more than the Xtt strains, though the populations were not significantly different between Xtu CO237 and Xtu ICMP11055, nor Xtt CO237 and Xtt UPB886. Between barley and wheat, Xtu populations reached similar levels, whereas Xtt populations were significantly greater in barley. Overall, we found that bacterial populations do not account for the increased severity of symptoms observed for Xtu CO237 and Xtt CO236, suggesting that virulence is determined by a factor outside of bacterial populations.

### Complete genome sequences of Xtt CO236 and Xtu CO237 reveal similarities between other Xtt and Xtu genomes

To investigate whether virulence of CO236 and CO237 is associated with major genomic rearrangements, we sequenced the two Colorado isolates using single-molecule long read sequencing (Oxford Nanopore Technology) (**Figure 2**). Nanopore sequencing yielded a total of 8.9 Gb of base-called reads for Xtt CO236, and 12 Gb for Xtu CO237. Filtong [30] was used to filter the reads, and a final total of 4.4 Gb for CO236 and 5.9 Gb for CO237 was used to generate whole-genome assemblies. With >100x coverage, the final genome assemblies were confirmed with CheckM [34] to be >99% complete. The draft Xtt CO236 and Xtu CO237 genome assemblies from the raw base called reads did not resolve the TALomes due to the formation of homopolymers in the sequence [17]. Therefore, we used Homopolish [32] to perform extra genome polishing after Flye-based genome assembly, and two polishing steps of the raw FastQ reads using Medaka 1.5.0 [15] to resolve the specific TALEs. The TALE classes were then annotated and assigned into their respective classes using AnnoTALE [21].

**Figure 2.**
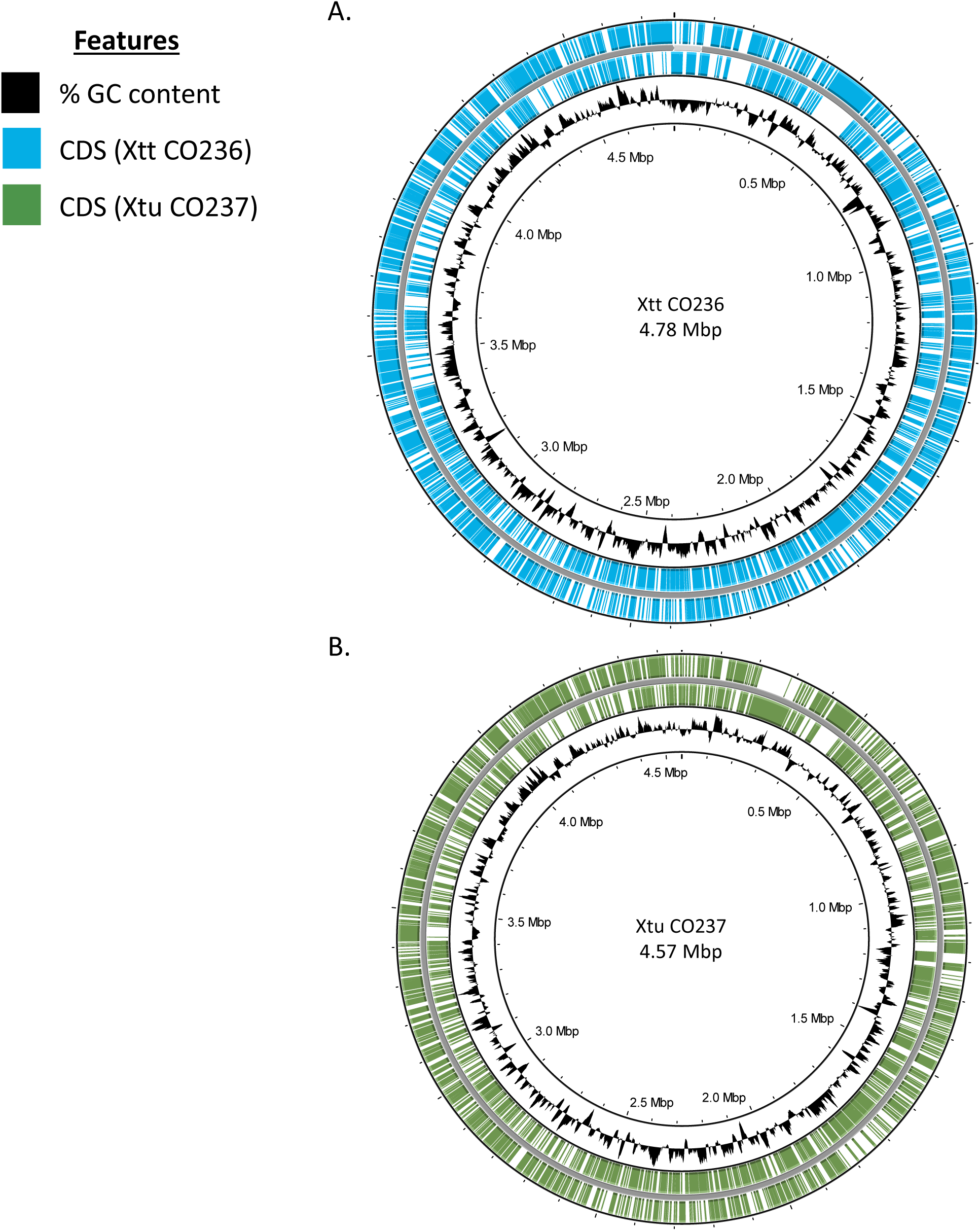
Genomic sequence maps of Colorado isolates A) CO236 (*X.t.* pv translucens) and B) CO237 *(X.t.* pv. undulosa). Genomic maps were generated *de novo* using Nanopore sequencing reads and Flye. GC: Guanine/Cytosine, CDS: Coding Sequence, Mbp: Mega-base pairs. Circular plots representing bacterial genomes were made using the Proksee webserver (https://proksee.ca).

The Xtt CO236 genome is 4,781,527bp with 3,767 genes predicted using a general prokaryotic pipeline [33], with an overall 67.86% GC content (**Table 1**). A small contig of 88,027 bp was also detected, and we predict that this is a large plasmid because there are no overlaps with the rest of the genome, low sequence similarity, and a significantly different GC Skew from the rest of the chromosome (GC content of 62.33%). This is consistent with other Xtt genomes, including UPB886, which also has a predicted plasmid [25]. Xtu CO237 is 4,567,484 bp base pairs with 3,680 genes predicted [33], and an overall GC content of 68.05% (**Table 1**). Similar to other Xtu strains, no plasmid was detected in Xtu CO237. Both of our recent isolates Xtt CO236 and Xtu CO237 have a relatively high GC content (> 65%) compared to other bacteria, which is consistent with other *Xanthomonas* species [35]. Some areas of the genome only had predicted coding sequence (CDS) for one strand (+/-). In areas of the genome where this was the case, there were several CDS predicted in one direction.

**Table 1.** Comparison of genomic features of Xtt CO236 and Xtu CO237 with other published Xtt and Xtu strains. Data for published isolates from ^1^Roman-Reyna et al., 2020; ^2^Shah et al., 2021; and ^3^Falahi Charkhabi et al., 2017 were collected from NCBI. TAL: Transcription Activator Like; T3: Type III.

|  | *X. translucens* pv. *translucens* |  |  | *X. translucens* pv. *undulosa* |  |  |
| --- | --- | --- | --- | --- | --- | --- |
| Strain | CO236 | Km8<sup>1</sup> | UPB886<sup>2</sup> | CO237 | ICMP1105<sup>3</sup> | XT4699<sup>3</sup> |
| Contigs | 2 | 1 | 2 | 1 | 1 | 1 |
| Bases | 4,781,527 bp | 4,792,950 bp | 4,674,364 bp | 4,567,484 bp | 4,761,583 bp | 4,561,137 bp |
| Genes | 3,767 | 3,873 | 3,736 | 3,680 | 3,656 | 3,528 |
| rRNA | 6 | 6 | 6 | 6 | 6 | 6 |
| tRNA | 53 | 53 | 53 | 54 | 54 | 54 |
| T3 Effectors | 34 | 34 | 36 | 31 | 32 | 32 |
| TAL effectors | 5 | 8 | 5 | 8 | 7 | 8 |

The Xtt CO236 genome is significantly larger than the Xtu CO237 genome (∼200kb difference), and Xtu CO237 is smaller than Xtu ICMP1105 (∼200kb difference). However, the total sizes of Xtu CO237 and Xtu XT4699, another highly-virulent United States isolate, are similar. Canonical type III effectors (T3Es), rRNA, and tRNA numbers were similar between Xtt CO237, Xtu CO236, and other Xtt and Xtu isolates (**Table 1**). The number of transcription activator-like (TAL) effectors was the same between Xtu CO237 and Xtu XT4699 with eight predicted, which is only one more predicted TAL effector than Xtu ICMP1105. Xtt CO236 and Xtt UPB886 both have five predicted TAL effectors, while Xtt Km8 has eight. Overall, Xtt CO236 and Xtu CO237 genomic patterns are similar to other Xtt and Xtu isolates. Therefore, major genomic changes do not obviously explain the differences in virulence between the Colorado isolates and other isolates.

### Whole-genome average nucleotide analysis of *X. translucens* genomes show groupings of closely-related sequences by pathovar

Next, we investigated whether genomic differences at the pathovar level could explain differences in virulence. All *X. translucens* genomic sequences deposited into NCBI by September 1, 2023 were compared for their average nucleotide identity (ANI) and grouped according to sequence similarity (**Supplemental Table S2**). **Figure 3** displays a heat map showing the comparative average nucleotide identity of each genomic sequence and associated group. The total counts of nucleotide identity % were obtained using PyANI [36], and further analyzed independently by seaborn’s clustermap function [37]. Strains were sorted into distinct pathovar groups, suggesting that genomic sequences among these clades are more similar to each other than other pathovars (**Supplemental Figure S2**). As expected, the Xtt CO236 genome was closely related to other strains classified as pathovar translucens, and Xtu CO237 was highly similar to other pathovar undulosa strains. Both isolates grouped with their respective pathovars. Within the largest group of deposited genomes, Xtu strains showed an overall similar genomic identity, and a subgroup of these strains had > 97% genomic identity between them (**Figure 3).** The strains annotated as pathovar cerealis also grouped closely together, suggesting that they are genomically distinct from Xtt, Xtu and other pathovars, which is in agreement with previous research [38, 39]. There was high sequence conservation at the chromosomal level between each of the pathovars, with the least similarity between the less-described graminis (‘grasses’) group, consisting of pathovars poae, phlei, phleiphratensis, arrhenatheri, and graminis, for which there are few full-genome sequences available. Two genomes annotated as pathovar hordei (UPB458 and UBP947) grouped with pathovar translucens. *X. translucens* pv. hordei was a previous taxonomic designation that has been reclassified as *X. translucens* pv. translucens [39, 40]. No apparent outliers with strong sequence differences appeared between pathovars.

**Figure 3.**
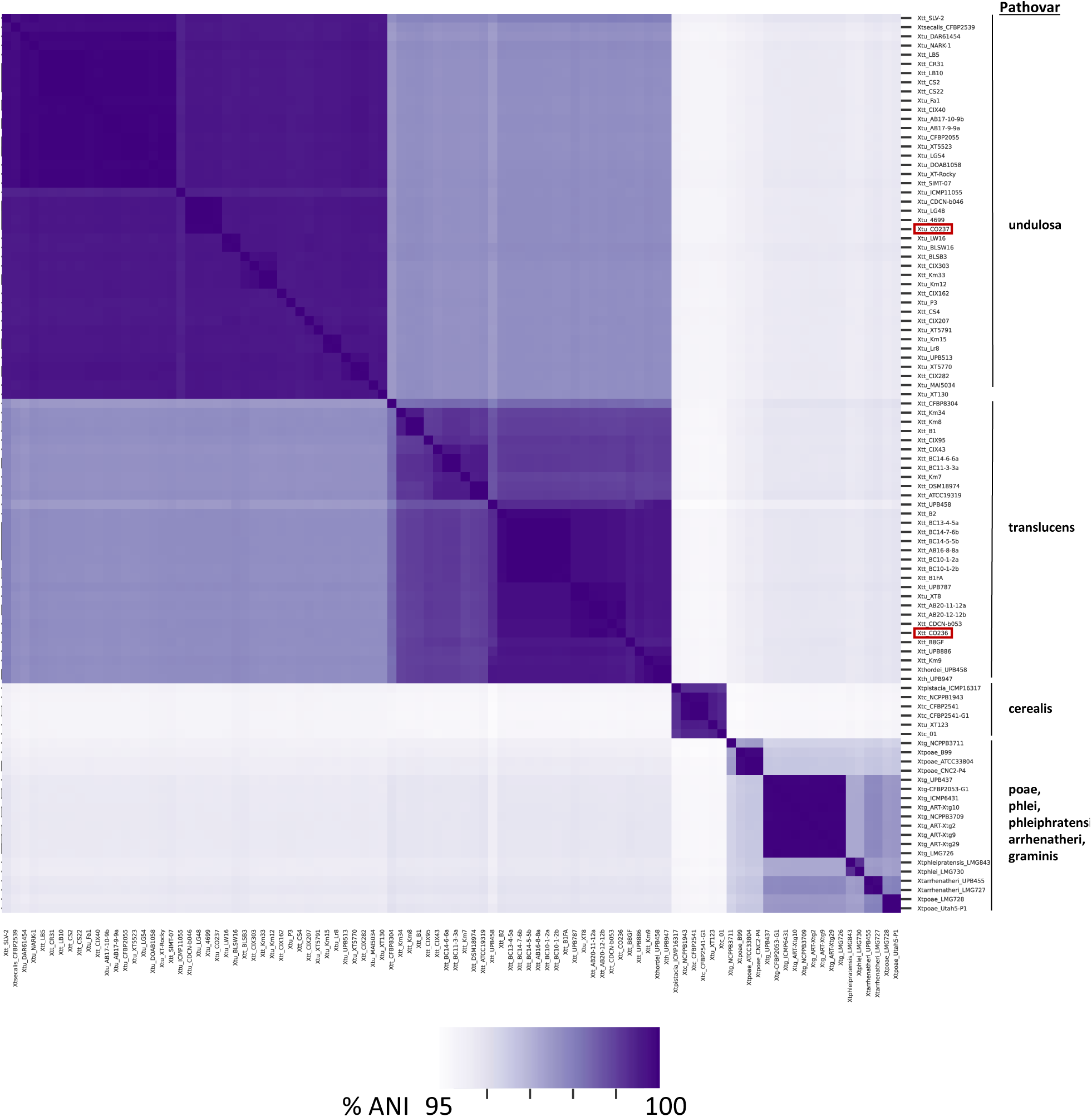
Average nucleotide identity (ANI) of *X. translucens* pathovars. Heatmap shows ANI between all *X. translucens* complete genomes available from NCBI (as of August 2023). Red boxes highlight isolates reported from the present study (Xtt CO236 and Xtu CO237). See Supplemental Table 1 for individual comparisons between strains.

### TALEs are sequence-diverse for Xtt CO236 and Xtu CO237

While the numbers of effectors (T3Es and TALEs) were similar between the Colorado isolates and the older strains, we hypothesized that differences in effector diversity may be correlated with virulence. We compared the TALEs, TALE classes, and RVDs between Xtt CO236, Xtu CO237, and other Xtt/Xtu isolates, and found that Xtt CO236 and Xtu CO237 encode diverse TALE classes (**Table 2**). Xtt CO236 encodes five TALEs, each from a different class (Tal DA, Tal CT, Tal CV, Tal JE, and Tal IY). These TALEs are located in three clusters on the genome, with TALE classes DA, CT, and IY located closely together on the chromosome, and TAL classes CT and CV being in separate and distinct genomic regions from the other TALEs (**Table 2** and **Figure 4**). Xtu CO237 encodes eight TALEs from eight different classes (Tal DD, Tal DA, Tal CZ, Tal DE, Tal CT, Tal DF, Tal DB, and Tal DC). These TALEs were also genomically clustered into four groups, with each cluster comprising of two TALEs from different classes (DD and DA, DB and CZ, DE and DF, and CT and DC). Interestingly, we found no overlap in the TALE classes between CO236 and CO237.

**Figure 4.**
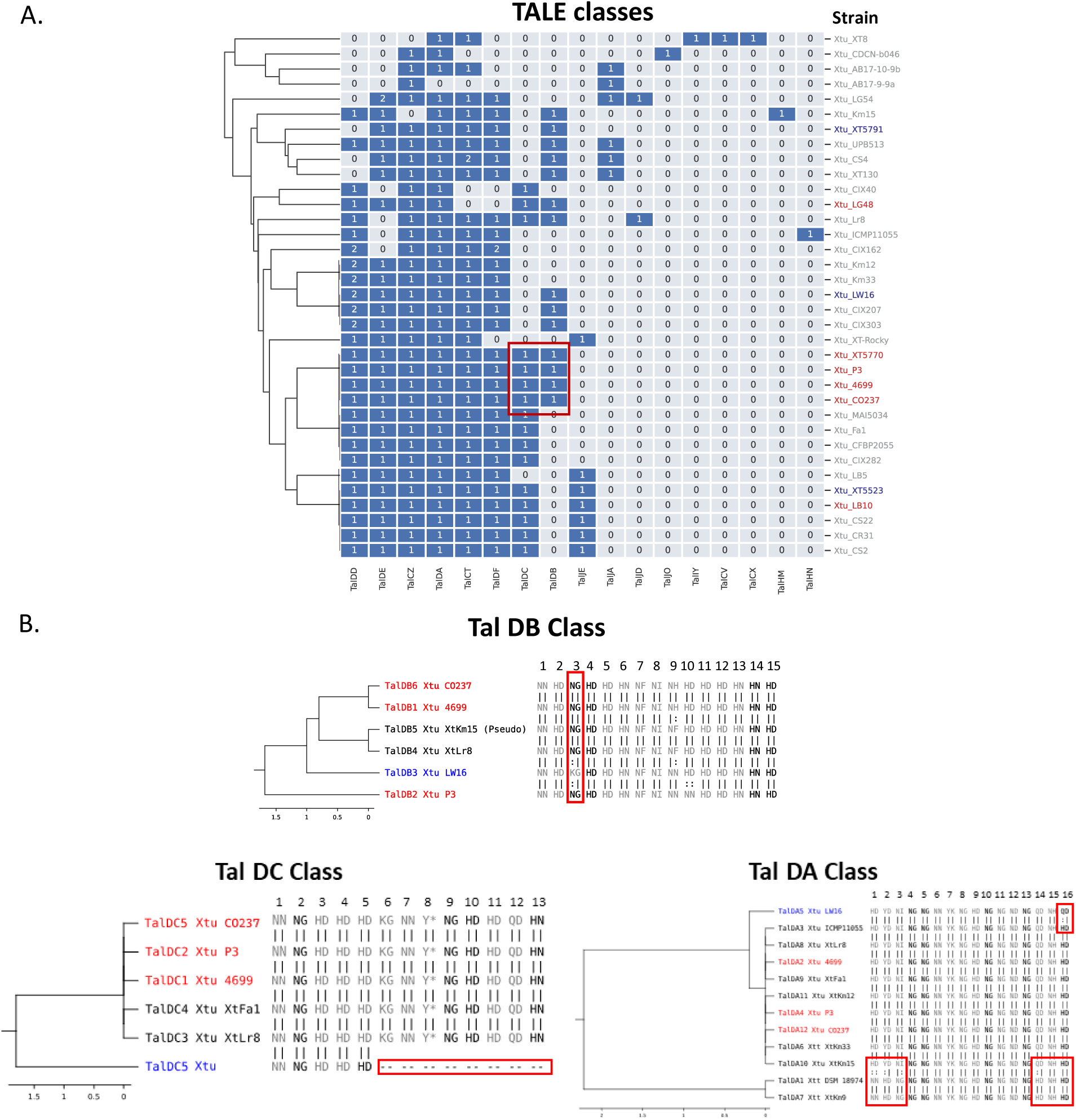
Effector analysis of Xtu isolates. A. TALE classes in published *X. translucens* genomes. Published Xtu strains without annotated TALEs were not included in the analysis. Numbers represent the number of copies of each TALE present in the genome. The TALE classes were annotated with AnnoTALE [21]. High virulence (red) and low virulence (blue) strains are indicated as published in [22]. TAL classes from high- and low-virulence strains produced by AnnoTALE. **B.** TALE class tree with assigned TALEs from Xtu CO237. Class assignment and trees were made with AnnoTALE. Number above represents the RVD repeat number.

**Table 2.**
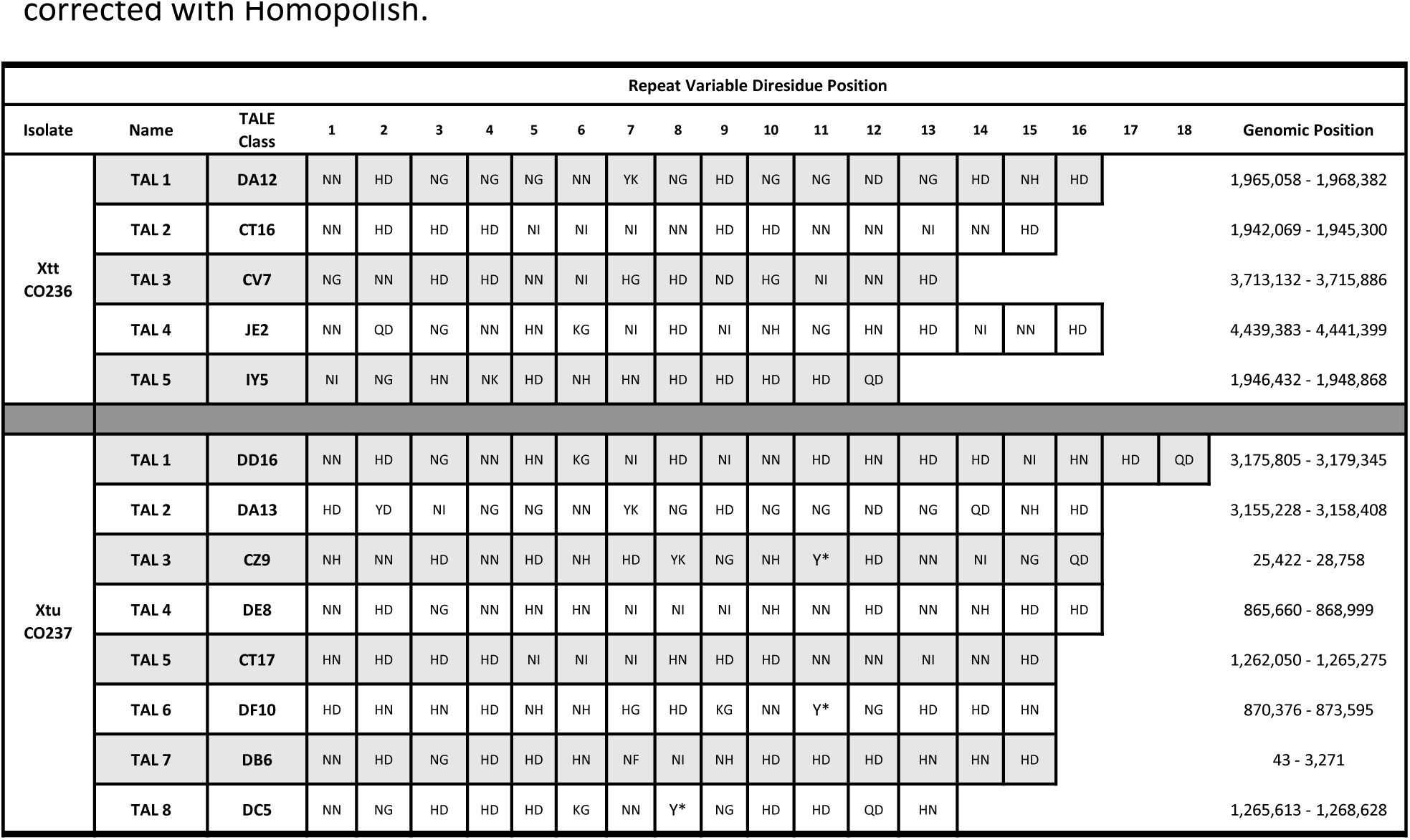
Analysis and comparison of TAL effectors present in Xtt CO236 and Xtt CO237. TALE classes, repeat-variable diresidues (RVDs), and genomic positions of the TALEs are listed. To identify all TALEs, the genome sequence was polished with Medaka and the TALomes were corrected with Homopolish.

Additionally, the repeat-variable diresidue (RVD) sequences of the individual TAL effectors were diverse and distinct from canonical *X. oryzae* TALEs. These included repeats YK, YD, Y* and QD, which are found in other *X. translucens* strains. The QD repeats were generally found at the end of the TAL effectors (Tal IY5, Tal DD16 and Tal CZ9), except for Tal DA13 and Tal DC5. Repeats YK, YD and Y* have not been described to bind to a specific nucleotide compared to other repeats that have different binding affinities to A, T, C and G [41].

### TALE classes correlate with *Xtu* virulence

Focusing on the Xtu-wheat pathosystem, we next explored effector diversity within available Xtu genomes and compared them to Xtu CO237. To first look at canonical, non-TALE type III effectors (T3Es), we generated a proteome database based on the predicted proteins of all of the published *X. translucens* genomes on NCBI (September 2023). A list of putative *Xanthomonas* effectors obtained from the *Xanthomonas* Type III Effector Database [42] was BLASTed against our custom proteome database (**Supplemental Table 3**). Based on a 75% protein homology threshold, Xtu strains were annotated with these predicted T3Es. A core set of 13 Xops (XopZ, XopX, XopV, XopQ, XopN, XopK, XopG, XopF2, XopB, XopAM, XopAF, XopAA and XopAD) was present in all of the Xtu strains analyzed (**Supplemental Figure S3).** A large portion of these strains (19/30) also carried XopJ5, while a lower proportion (4/30) had XopL, and only two strains possessed XopH1. Overall, the non-TALE T3Es were highly conserved among Xtu strains.

Next, we compared the TALE classes between the same set of Xtu strains. A subset of 5 strains (Xtu UPB513, NARK-1, DOAB1058, DAR61454 and BLSW16) did not have annotated TALEs and were not included in the dataset. Information about known, relative virulence of several strains, previously published [22], was overlayed with the presence/absence of TALE class to determine if there was a correlation with virulence (**Figure 4A**). Interestingly, we found that the high virulence strains (Xtu P3, XT5770, and 4699) and CO237 had similar TALE repertoires that were distinct from the strains classified with low virulence, with TalDC and TalDB primarily being present in the high virulence strains. However, two low virulence strains, Xtu XT5791 and XT5523, had both TalDC and TalDB classes, and LW16, another low-virulence strain, had TalDB. To further explore if this class was unique to highly virulent strains, we analyzed the RVD sequence of this class of TALEs and generated class trees to represent RVD similarity. We found that the RVD sequences of the high virulence strains were more similar than to the low virulence strains (**Figure 4B**). In particular, the low virulence strain LW16 has a TalDB TALE with a polymorphism in the third RVD repeat (NG ➔ KG). To explore whether this polymorphism could lead to changes in the predicted protein folding, the central repeat region (CRR) of the Xtu CO237 and LW16 TalDBs were computationally compared to each other using Alphafold 2 [43] (**Supplemental Figure S4**). We observed no significant differences in the CRR structure. While the effect of these repeats on the binding of the effector binding element (EBE) sequence is not clear, it is a conserved repeat in several TALEs from the Tal DB class (**Figure 4B**). Another low virulence strain that shared a TALE class with highly virulent group was Xtu XT5523. While TalDC is present mainly for high virulence strains (**Figure 4A**), in the low virulence strain Xtu XT5523 there is a major deletion that leads to only five of thirteen RVD repeats being present, whereas high virulence strains have a similar RVD sequence (Xtu CO237, P3, and 4699) (**Figure 4B**). The TalDA class is present in most of the available genomes (24/30) and is highly conserved in RVD sequence. The low virulence strain LW16 has only a single polymorphism in the last RVD repeat (HD ➔ QD). Other TAL effector classes with highly conserved RVD repeats can be found in **Supplemental Figure S5**.

Next, we asked whether there was a correlation between the location of the TAL effector within the genome and pathogen virulence. Because we observed that the TAL effector classes were clustered across the genome for the Colorado isolates (**Table 2**), we investigated whether this was true for other high virulence strains. We chose three high virulence strains and five strains of unknown virulence (not described in [22]) and mapped the TALE classes across the genome (**Figure 5A**). We found that all eight strains had similar clustering as Xtu CO237 for classes TalDA/DD and TalDC/CT, but CO237 was distinct in its clustering for TalDE/DF and TalDB/CZ. In CO237, TalDE/DF and TalDB/CZ exist in the same clusters. These clusters are broken in the other strains, such that TalDE and TalDF are physically separated, along with the separation of TalDB and TalCZ. Additionally, a translocation of these four TALE classes physically moves the position of the TALEs relative to the other strains. In four of the strains (CFBP2055, ICMP1105, Fa1, and Km12), the physical position of TalDC/CT changes relative to the other TALEs, though the cluster does not break. We then looked at the gene orientation of the TALEs in CO237 and compared it to the two most closely related Xtu isolates, 4699 and P3 (**Figure 5B**). We found that not only has the order of the TALEs changed, but the orientation of TalCZ is also reversed. Overall, the CO237 TAL effector classes are positioned in ways unique from other Xtu strains.

**Figure 5.**
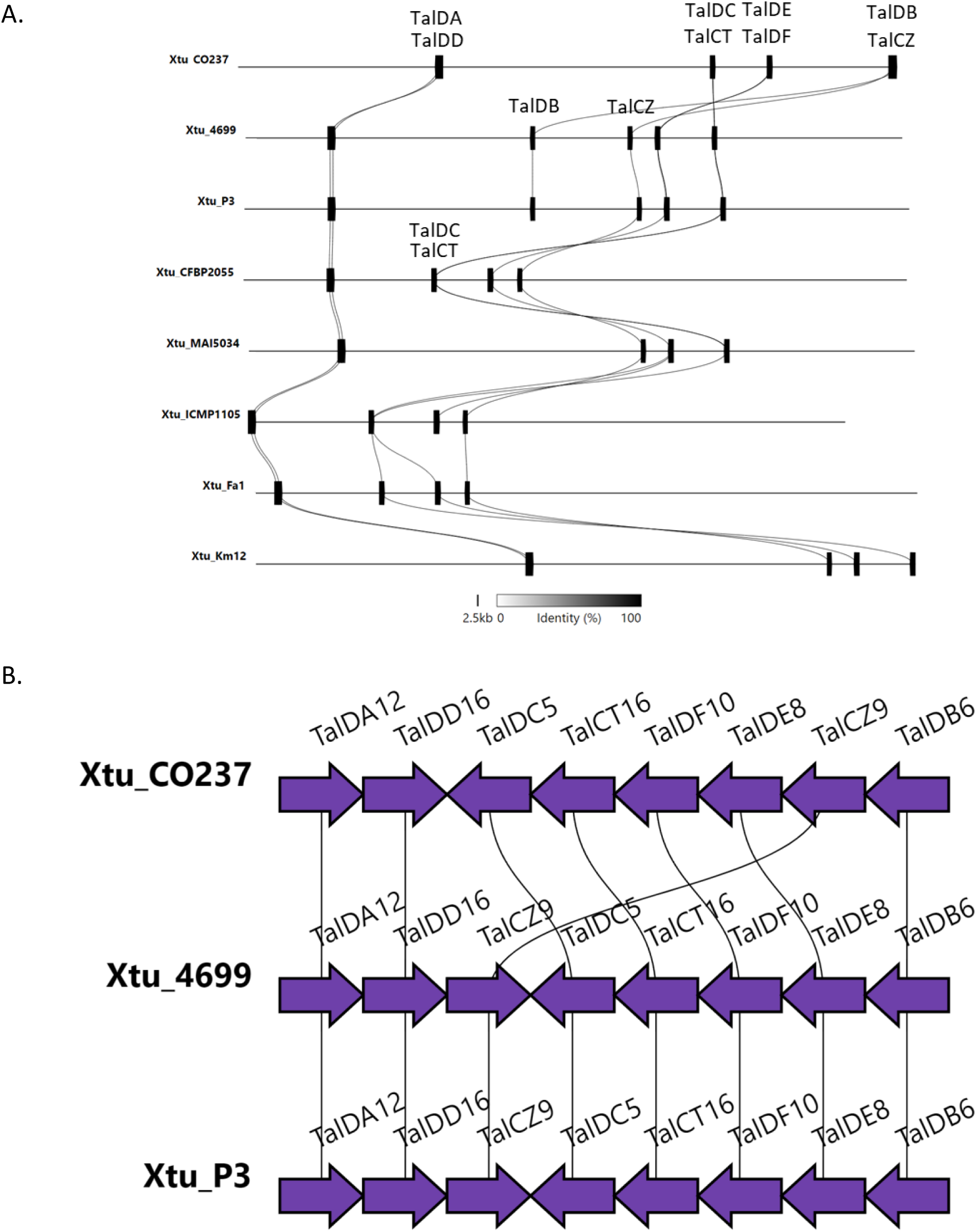
TALE alignment in complete genomes of *X. translucens* pv. undulosa. A. Complete genome maps of eight Xtu strains with mapped TALE classes across the chromosome. **B.** Simplified version of the TALE repertoire of three highly virulent Xtu genomes, showing differences in TALE gene orientation across genomic positions within the bacterial chromosome. Figures **A** and **B** were made using Clinker.

## DISCUSSION

Emerging and re-emerging plant pathogens pose a major threat to global agriculture and food security, especially in the context of climate change, and disease outbreaks have significant impacts on crop breeding programs which typically take years or decades to incorporate new disease resistance [3, 44–46]. Bacterial leaf streak of cereals, caused by *X. translucens*, is a re-emerging disease in that it is resurging into a problematic disease that will likely impact the wheat production landscape worldwide [3].

We characterized two recently collected *X. translucens* isolates from Colorado (Xtt CO236 and Xtu CO237) with increased virulence compared to previously described isolates (**Figure 1**). While the symptoms of the Colorado isolates were more severe than older isolates, the bacterial population sizes did not correlate with increased virulence, suggesting that a virulence factor within the pathogen aids it in increased aggressiveness. To address whether genomic changes could impact virulence in the Colorado isolates, we sequenced the genomes of Xtt CO236 and Xtu CO237 using Oxford Nanopore Technology, and compared the genomes with previously described Xtt and Xtu genomes (**Figure 2**). Xtt CO236 was very similar to other Xtt isolates, and Xtu CO237 was very similar to other Xtu isolates at the genomic level. In *Xanthomonas*, the expansion to new hosts and the globalization of crops has caused this genus to be very diverse in its genomic features. However, our results (**Figure 3**) show that *X. translucens* strains have over 95% similarity between all genomes, suggesting a conserved genomic repertoire of virulence factors and chromosomal structure. The highly conserved genomic structure of *X. translucens* strains suggest that there is a strong selective pressure to maintain the repertoire of virulence factors and genomic architecture, which could confer a fitness advantage to the pathogen in their host plants. A potential explanation could be that *X. translucens* has evolved to become highly specialized for infecting cereal crops, leading to less genetic diversity within strains. Moreover, there is a clear distinction of DNA similarity between strains from different pathovar classifications. This supports our previous hypothesis that *X. translucens* is likely to have evolved high specialization in their respective hosts, showing closely related genomes among strains from the same pathovar.

Alongside management of this disease, there have been different reports supporting reclassification of strains from this species, instead using the LINgroups system [7, 47]. This new classification system clearly separates strains from pathovars translucens and undulosa, although it is a genetic based algorithm, rather than the initial descriptions of pathovar [5]. Our results with whole-genome average nucleotide identity analysis support these data and show the importance of using whole-genome data for identifying and characterizing virulent strains that could emerge or become more prevalent. The development of new and inexpensive technologies for large collections of *Xanthomonas* species can further support the establishment of standards for bacterial identification which could aid in plant protection and rapid diagnostic.

Like some other Xtt isolates, Xtt CO236 has a predicted plasmid. The role of this plasmid has not been tested, but typically, plasmids can aid pathogens in rapid adaptation to a host’s environment [48]. Because *X. translucens* is re-emerging under the selection pressure of climate change and new cereal varieties, the plasmid may aid Xtt in host adaptation or virulence. The genome sequence of Xtt is more diverse than that of Xtu (**Figure 3**), which may suggest that Xtt is more rapidly evolving than Xtu, even though both pathovars are likely under similar selective pressures since barley and wheat are closely related and grow in similar environments. Pathogen infection lifestyle may play a role in this selection, since Xtt is generally vascular and Xtu is generally apoplastic. While Xtt strains are very similar to each other on a genomic level, strain virulence factors are globally distributed [49]. There is largely no overlap between the TALE class repertoire between different pathovars, except for classes TalCT and TalDA (**Supplemental Figure 2**). This shared class would suggest that there are similar targets in their corresponding hosts (wheat and barley), as these classes will target similar promoter sequences within the genome. However, Xtt strains have a wider repertoire of classes not shared with any Xtu strain, including TalCX, TalCV, and TalIY and Tal JA. This suggests that there have been clear distinctions of the pathogen adaptability to the host. It is possible that newer cereal varieties have greater selective pressures within the plant vasculature, as they have been selected for increased yield, quality, drought tolerance, and other disease resistance traits.

Bacterial virulence and aggression are typically linked to the expression of different effector repertoires, which can increase virulence by suppressing the plant immune system, making nutrients more available to the pathogen, or changing the plant environment to better support population growth, making virulence factors indispensable for disease development [50, 51]. TALEs are one example of a class of virulence effectors, which typically suppress plant resistance genes and/or activate susceptibility genes by modulating plant gene expression through plant promoters [52]. While there is an energy cost associated with gene expression, diverse effector repertoires can aid pathogens in adaptation to different plant environments and may be expressed only under the appropriate environmental conditions. Xtt CO236 and Xtu CO237 both have diverse effector repertoires with different repeat variable di-residues (RVDs) within the TALE central repeat region (CRR), which is responsible for host promoter binding (**Table 2**). The non-canonical repeats found in the CRR, including YD, YK, Y*, and QD, have been found in other *X. translucens* strains [22, 26]. Although the contribution of these repeats to TALE efficiency or effectiveness has not been determined, it is hypothesized that the repeats could enhance the upregulation of their targets [26]. Moreover, upregulation of these TALEs could enhance virulence in emerging Xtu strains compared to historical isolates. The RVD diversity between Xtt CO236 and Xtu CO237 may suggest that these isolates are, and/or have been, under constant selection pressure to adapt to their host’s environment. Looking at the TALE classes, there is no overlap between Xtt CO236 and Xtu CO237, which may also hint towards TALEs having impacts on host range and/or pathogen lifestyle. The mechanisms by which *Xanthomonas* species can upregulate the expression of virulence factors are quite diverse, including quorum-sensing signaling, cyclic di-GMP and two-component systems, regulation of *hrp* genes, and post-transcriptional regulation [53]. Determining the factors that make strains highly virulent could help us in developing strategies to manage virulence factors, increase plant resistance, and decrease yield losses caused by BLS.

We focused on the wheat-Xtu pathosystem because it is less understood than barley- or wheat-Xtt, and in Colorado, wheat is grown at larger acreage than barley. Additionally, we were interested as to how a pathogen that seems quite genomically conserved across the species could cause such drastic differences in symptom development across strains. Only a few TALEs have been previously described for Xtu [22, 26], and little has been described for other virulence factors [3]. Previous studies have looked into the genomic differences across *X. translucens* strains, which revealed a fairly conserved set of canonical T3Es, but diverse TALE repertoires [7, 49]. Similarly, we found that most Xtu strains had a conserved repertoire of twenty-three T3Es (**Figure 4A**). While T3Es were highly conserved among Xtu, there were striking differences across the TALEs. Two effector classes, Tal DC and Tal DB, were associated with more virulent strains, including Xtu CO237 (**Figure 4**). Two highly virulent strains (Xtu LG48 and LB10) did not cluster together with the other virulent strains in the effector analysis. This could be because they were not complete genomes, and putative TALEs are missing from the assembly. Another strain that had these classes (Tal DB and Tal DC) was Xtu Lr8, however, the relative isolate virulence is unknown. While a few low virulence strains encoded these two TALE classes, we found polymorphisms in the RVD domains of the low virulence strains compared to high virulence strains in both TalDC and TalDB classes, and this does not appear to be caused by major protein folding changes within the CRR (**Supplemental Figure 4**). Perhaps, we will find that binding affinity to the promoter is more influential than protein folding for TALEs. [22]. We also found that the positions of the TALEs within the strain genomes were largely conserved for seven Xtu strains, whereas major genomic shifts occurred in Xtu CO237 TALEs (**Figure 5**). This includes clustering of two groups in CO237, TalDE/DF and TalDB/CZ, which do not cluster together in other strains. These two classes are also rearranged in order on the chromosome, and TalCZ is also in the reverse orientation in CO237 compared to the seven other strains, as well as clustered next to TalDB. This major chromosomal rearrangement in CO237 may provide clues as to how this new isolate evolved and may hint at molecular mechanisms behind Xtu virulence.

Overall, we present genomic data from two highly virulent isolates collected from Colorado that suggest that virulence factors, including TALEs, may play an important role in increased isolate virulence compared to older strains. Genomic hallmarks show evidence of evolution of the genomes to fit pathogen host range and biological lifestyle, especially in the context of virulence factors, and may provide clues as to how future pathogens may emerge and/or increase in their aggressiveness in managed crops and ecosystems.

## Author contributions

*Diego E. Gutierrez-Castillo*: Methodology, Software, Validation, Formal Analysis, Investigation, Data Curation, Writing-Original Draft, Writing-Review and Editing, Visualization. *Emma Barrett:* Methodology, Validation, Investigation, Data Curation. *Robyn Roberts:* Conceptualization, Methodology, Formal Analysis, Resources, Writing-Original Draft, Writing-Review and Editing, Supervision, Project Administration, Funding Acquisition.

## Acknowledgements

We would like to thank Libby Swanson and Matt West for technical assistance with the project, Tammy Brenner and Paul Freebury for greenhouse assistance, Jan Leach and Emily Luna for gifting strains and bacterial resources, and Jan Leach for helpful comments on the manuscript. Kirk Broders and Tessa Albrecht collected the original CO236 and CO237 isolates in 2018.

This work utilized the RMACC Summit supercomputer, which is supported by the National Science Foundation (awards ACI-1532235 and ACI-1532236), the University of Colorado Boulder, and Colorado State University. The Summit supercomputer is a joint effort of the University of Colorado Boulder and Colorado State University.

## Supplemental Figures

**Supplemental Figure S1.**
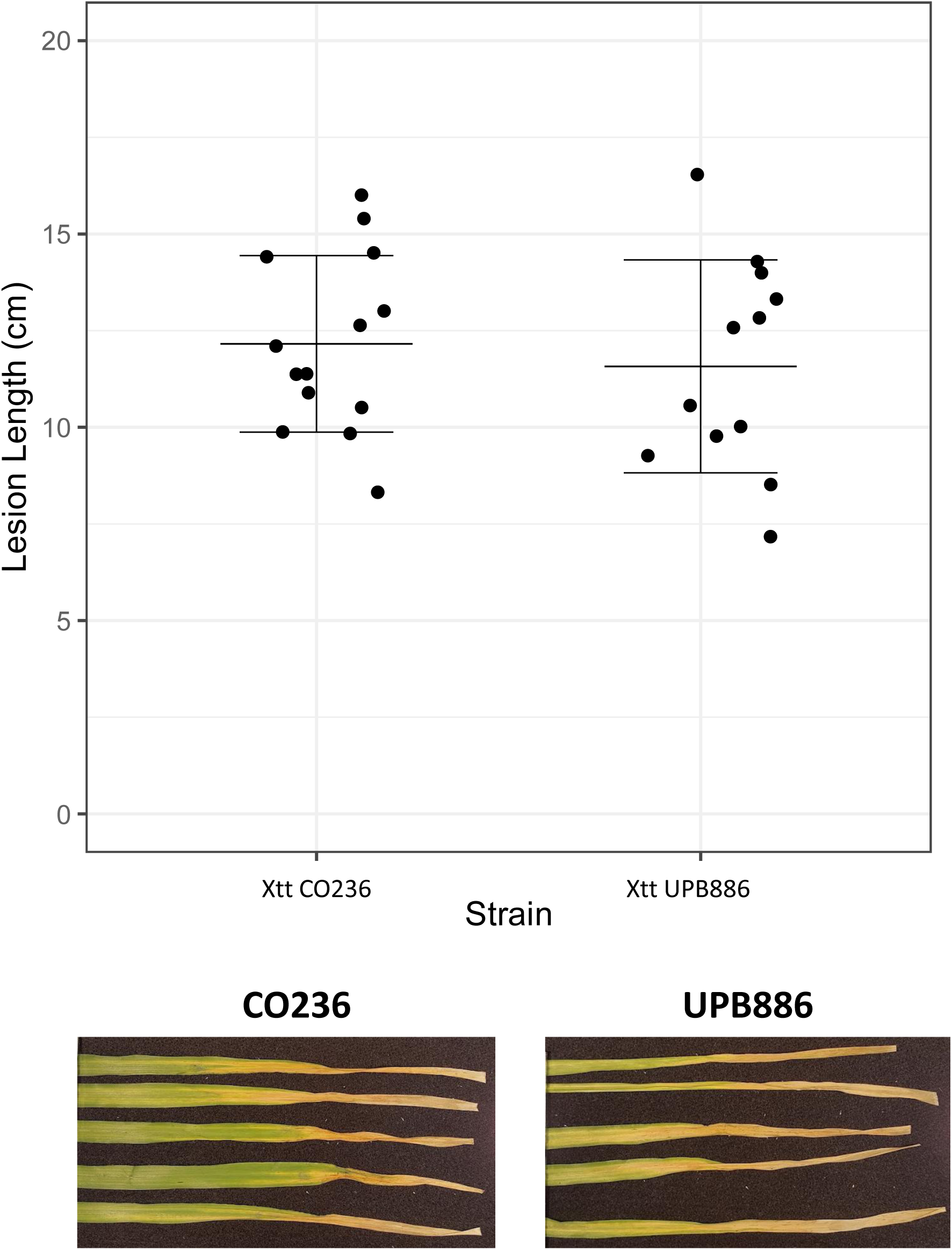
Lesion lengths from clip inoculations of CO236 and UPB886 in barley (var. Morex) leaves. Lesions were measured 14 days post clip-inoculation. No significant difference was found between Colorado isolate and UPB886. Significance values determined by Wilcoxon test in R software (p value = 0.55, n=12).

**Supplemental Figure S2.**
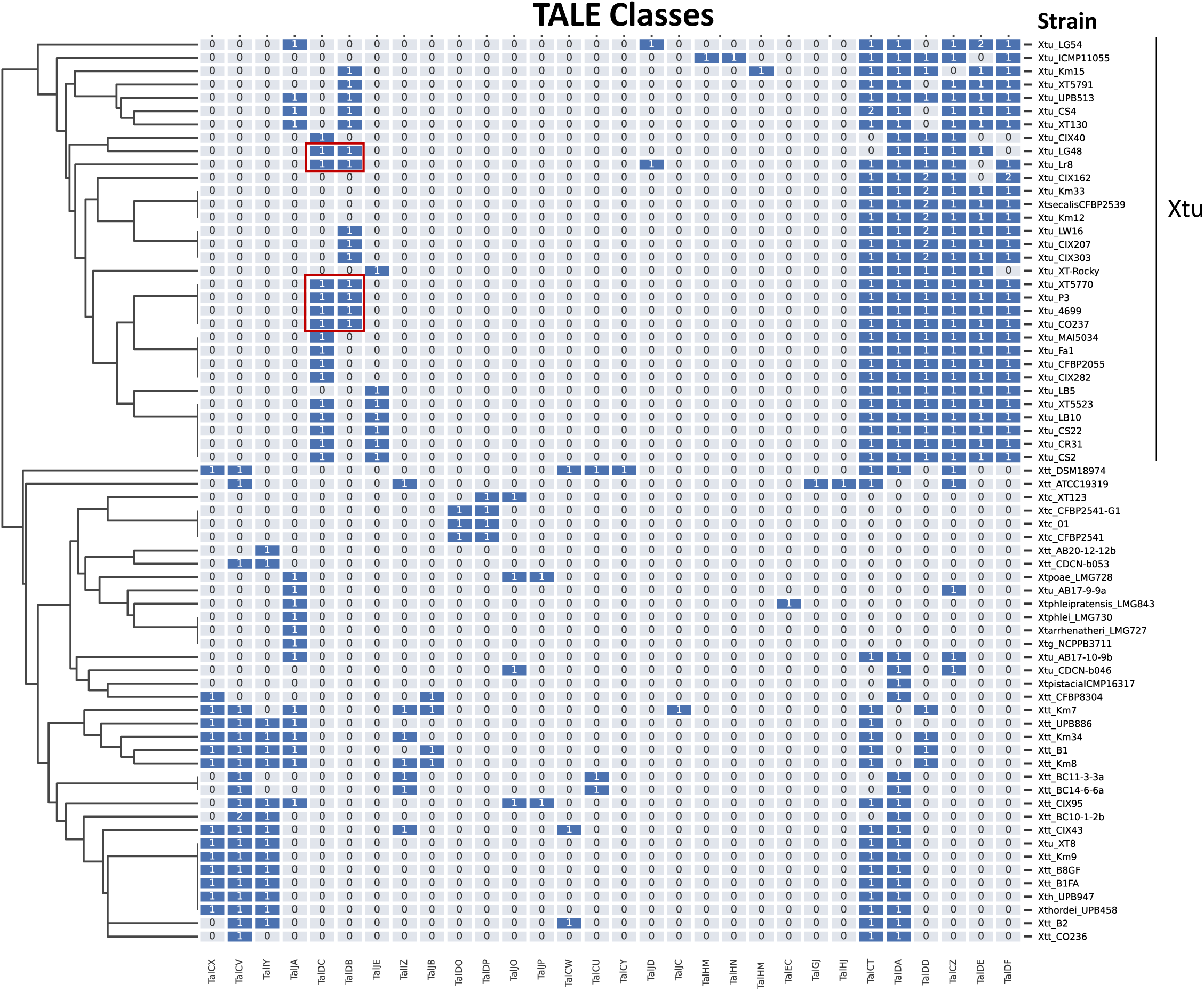
Effector analysis of all *X. translucens* isolates. TALE classes in published *X. translucens* genomes. Numbers represent the number of copies of each TALE present in the genome. The TALE classes were annotated with AnnoTALE (Grau et. al, 2016).

**Supplemental Figure S3.**
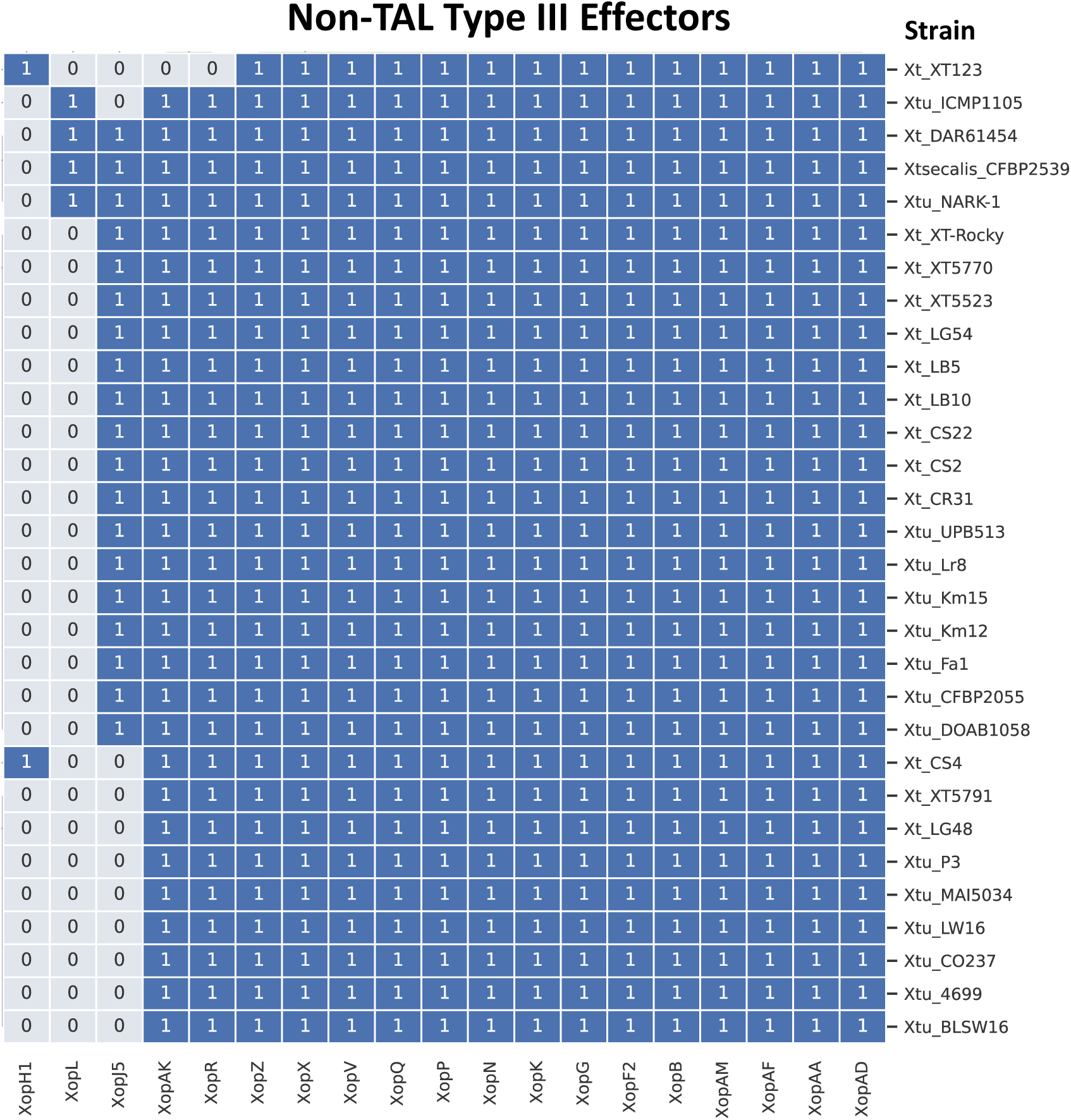
Non-TAL Type III effector homologs were determined using known *Xanthomonas* effectors as the query (obtained from: http://www.biopred.net/xanthomonas/t3e.html) and conducting a BlastP search in published *X. translucens* genomes. Numbers represent the number of copies of each effector in the genome.

**Supplemental Figure S4.**
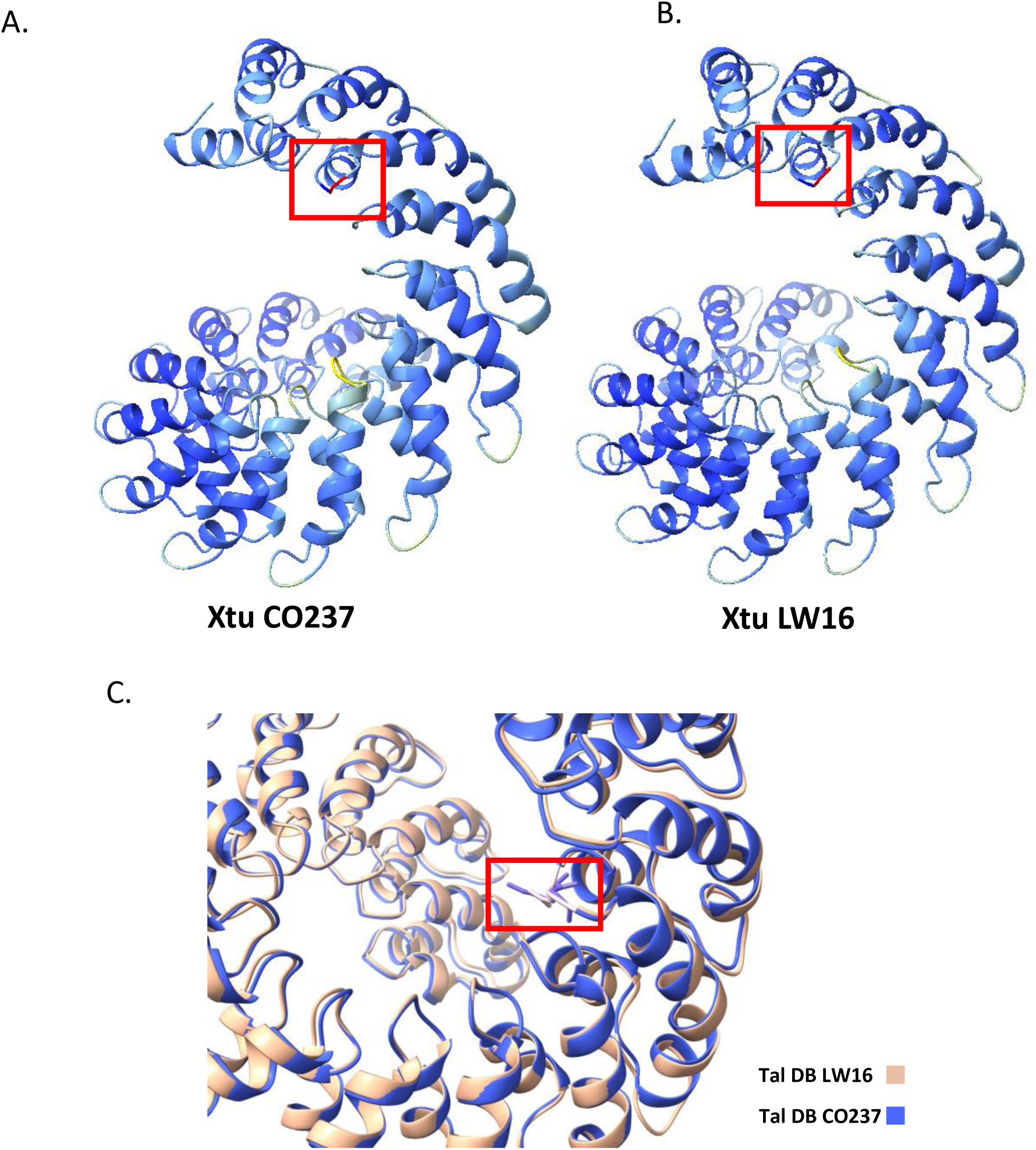
Protein structure prediction of the Central Repeat Region (CRR) of TalDB in Xtu CO237 (A) and Xtu LW16 (B). Alphafold2 was used to predict the protein structure of the CRR of two TALEs. Red boxes highlight the mutation in the third RVD (NG −> KG) of the TalDB class of the low virulence strain LW16. **C.** The overlay of both protein structures shows no apparent difference of the folding between the CRRs.

**Supplemental Figure S5.**
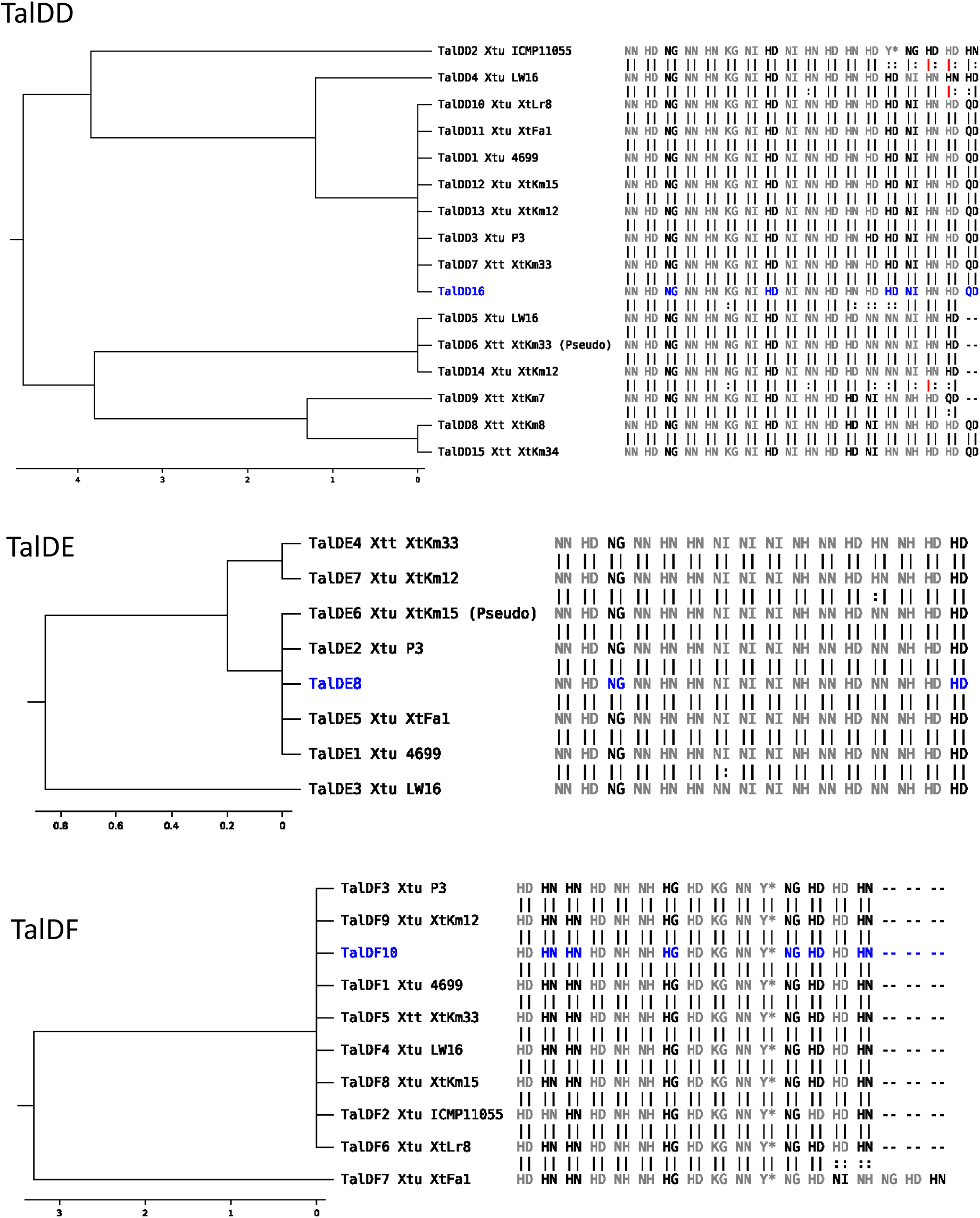
TALE classes in *X. translucens* pv. undulosa. TALE classes assigned by AnnoTALE. Xtu CO237 TALEs are highlighted in blue in each tree class (continued on next page).

**Supplemental Figure S5.**
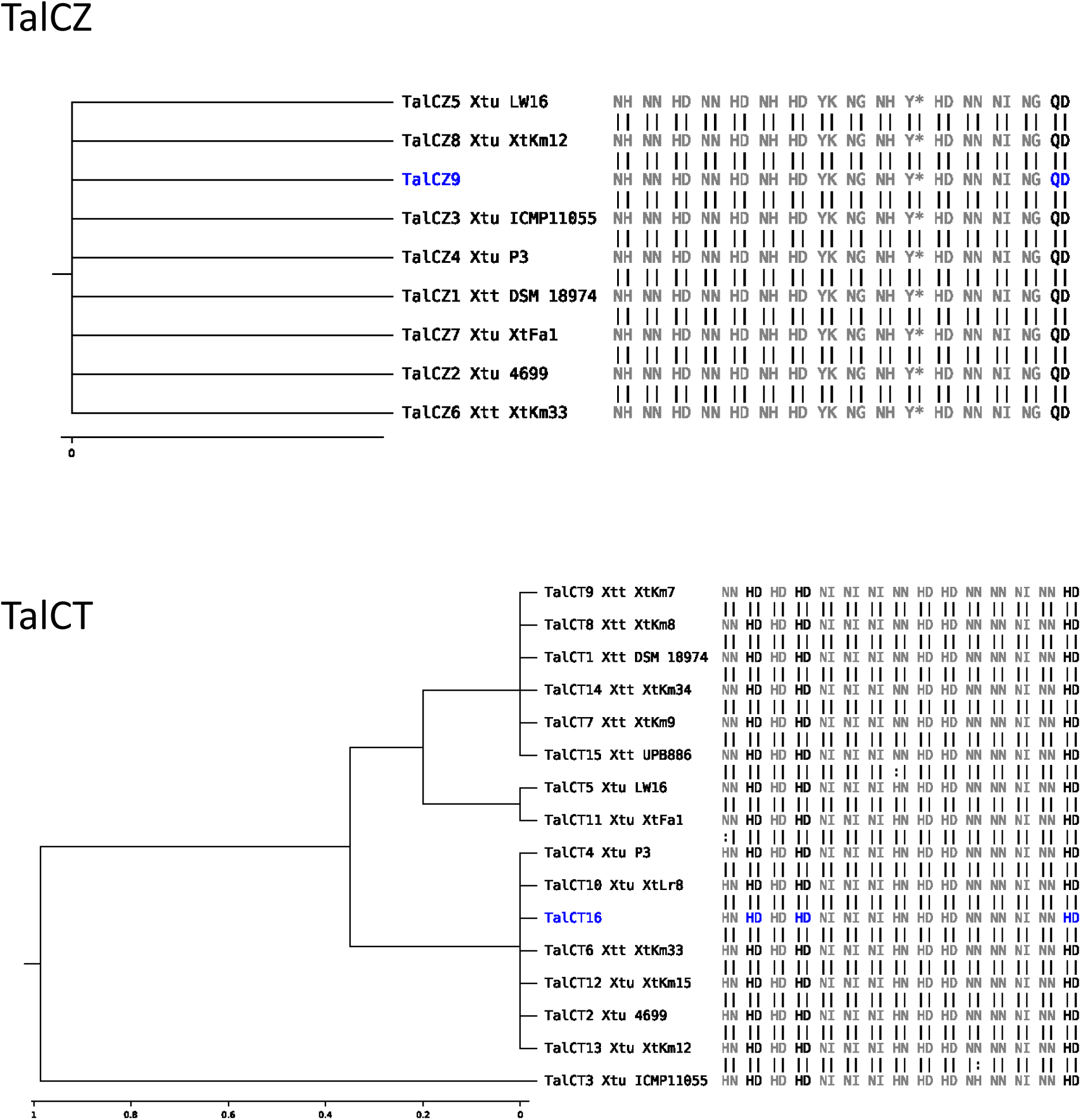
TALE classes in *X. translucens* pv. undulosa. TALE classes assigned by AnnoTALE. Xtu CO237 TALEs are highlighted in blue in each tree class (continued from previous page).

